# Dietary fatty acids modulate the endocannabinoid system in rat submandibular gland

**DOI:** 10.1101/2022.04.25.489419

**Authors:** César Nicolás Combina Herrera, Mariana Blanco, Gastón Repossi, Jorge Darío Escandriolo Nackauzi, Adriana Beatriz Actis

**Author notes:** **Corresponding author** Adriana Beatriz Actis, Instituto de Investigaciones en Ciencias de la Salud, Consejo Nacional de Investigaciones Científicas y Técnicas, CONICET, FCM, Universidad Nacional de Córdoba. Bv. de la Reforma y Enfermera Gordillo, Ciudad Universitaria, 5000 Córdoba, Argentina. The last co-authorship is shared between Jorge Darío Escandriolo Nackauzi and Adriana Beatriz Actis.

## Abstract

The aim of this study was to analyze the influence of dietary fatty acids on endocannabinoid system components in rat submandibular gland. 15 male Wistar rats were fed on commercial diet until the 8th week of life. They were then divided into three dietary groups: Control, continuing with chow diet, and two experimental groups receiving for 30 days a diet containing corn oil or chia oil as lipid source (7%). At that time, animals were sacrificed and salivary glands obtained. Anandamide and 2-arachidonylglycerol endogenous ligands (high performance liquid chromatography) and cannabinoid receptors CB1 and CB2 (immunofluorescence) were quantified. Fatty acid amide hydrolase enzyme activity was analyzed (spectrophotometry). Kruskal Wallis test was applied. 2-Arachidonylglycerol was higher in Corn oil group than in Control and Chia oil groups. The greatest CB1 and the lowest CB2 receptor positively-marked area percentage were found in Corn oil and in Chia oil, respectively. Fatty acid amide hydrolase enzyme activity was higher in Corn oil than in Chia oil. 18:2 n-6 (Corn oil) and 18:3 n-3 (Chia oil) dietary fatty acids modulate endocannabinoid system components in rat submandibular gland, what would have influence on salivary secretion. Dietary n-3 fatty acid could be useful in salivary dysfunctions.

## Introduction

Salivary physiology may be affected by various factors such as age, circadian rhythms and diet, which modify saliva chemical composition and secretion [1,2]. Alam and Shi [3] reported that dietary-induced essential fatty acid deficiency produces changes in their composition of rat major salivary glands, which are reflected in salivary total lipids.

Previously, we found that fatty acid composition of rat submandibular glands, as well as the salivary flow are modulated by their dietary intake and by the time elapsed since their consumption. Stimulated submandibular saliva was higher in animals fed on n-3 fatty acid-rich diet (chia oil) as compared to those fed on n-6 and n-9 fatty acid-rich diets (corn and olive oils, respectively). A positive correlation between this functional finding and the fatty acid glandular profile was also observed [4].

It is known that endocannabinoids have influence on salivary secretion [5,6]. These bioactive fatty acid-derived molecules are synthesized on demand by cells, not stored and rapidly degraded after their action. Endocannabinoids are involved in a wide range of physiological functions like cognition and memory, control of appetite and energy metabolism, modulation of immune responses, reproduction and fertility, among others [7]. They mainly comprise arachidonylethanolamide or anandamide and 2-arachidonylglycerol endogenous ligands [8], which are part of a complex system also composed by cannabinoid receptors (CB1 and CB2) and all the enzymes involved in their synthesis and degradation: N-acyl phosphatidylethanolamine phospholipase D, monoacylglycerol lipase, diacylglycerol lipase and fatty acid amide hydrolase [9,10]. Endocannabinoids are synthesized from membrane lipids due to an increase in intracellular calcium levels. Once released they can be recaptured through a complex mechanism of high affinity membrane transport [11].

Dietary n-6 fatty acid intake is the main source of endocannabinoids. Nutritional changes may affect their levels since the metabolic pathway responsible for the synthesis of anandamide and 2-arachidonylglycerol from arachidonic acid is also involved in the synthesis of eicosapentaenoyl ethanolamine and docosahexaenoyl ethanolamine from n-3 eicosapentaenoic and docosahexaenoic acids, which compete with anandamide and 2-arachidonylglycerol for the same receptors (CB1 and CB2). High levels of n-3 fatty acid-derived endocannabinoids trigger physiological responses that are opposite to those of anandamide and 2-arachidonylglycerol [7]. Thus, variations in dietary consumption modifying the n-6/n-3 fatty acid ratio have impact on the incorporation of their derivatives and on the endocannabinoid/ethanolamine synthesis [12,13].

Concerning the influence of endocannabinoids on salivary secretion, anandamide decreases it through the activation of CB1 and CB2 cannabinoid receptors located in acini, ducts and nerve endings of submandibular gland [5,14]. In addition, it was observed that bacteria lipopolysaccharide injected intraperitoneally inhibits the salivary secretion by increasing the production of prostaglandins and endocannabinoids and that anandamide acts on hypothalamic CB1 receptors inhibiting the salivary secretion by attenuating the parasympathetic neurotransmission to the submandibular gland [6].

The modulatory role of the endocannabinoid system on the submandibular gland function has been described but not in relation to dietary fatty acid intake. The aim of this study was to analyze the influence of dietary fatty acids on the endocannabinoid system components - endogenous ligands, cannabinoid receptors and fatty acid amide hydrolase enzyme activity-in rat submandibular gland.

## Materials & methods

### Animals and diets

Fifteen healthy adult male Wistar rats obtained from the Mercedes and Martín Ferreyra Research Institute and kept at the Institute of Health Sciences Research (Instituto de Investigaciones en Ciencias de la Salud, INICSA), CONICET-UNC, Córdoba, Argentina, were employed. Each rat (body weight = 300 ± 50 g) was housed in a separate cage under standard temperature conditions (23 ± 2 °C), with 12-hour light-dark cycles. Animals had access to food (commercial or laboratory diet) and water *ad libitum*. Body weight (g) and dietary intake (g/day) were registered (weekly and daily, respectively). The animal handling was done according to the international guidelines for the care and use of laboratory animals [15]. The study protocol was previously approved by the Institutional Ethics Committee on Health Research (CIEIS) of the Dental College, National University of Córdoba, Argentina (Protocol # 12/2013).

After weaning (3 weeks), all animals received a standard commercial chow (Balanceado Asociación Cooperativa Gilardoni) until the 8^th^ week of life. They were then divided into three dietary groups: Control -continuing on chow diet- and two experimental groups feeding on a semi-soft laboratory-made diet with corn oil or chia oil as lipid source (7%) for 30 days (Corn oil and Chia oil groups, respectively). Diets contained the essential nutrients required for laboratory animals [16] and were adjusted to provide similar calories.

Fatty acid composition of diets was analyzed by gas chromatography. Lipids from Control, Corn and Chia oil diets were extracted by means of chloroform-methanol 2:1 according to Folch’s method [17]. Fatty acids were then methylated by sodium methoxide [18] and dried into a nitrogen atmosphere. Fatty acid methyl esters were analyzed by gas chromatography by means of a Shimadzu 2014 chromatograph equipped with a flame ionization detector.

Temperature of both injector and detector was 250 °C, and nitrogen was the carrier gas. Fatty acid methyl esters were analyzed by using a 100 m×0.25 mm×0.2 mm film thickness SP-Sil 88 capillary column (Varian). The results are expressed as percentages of total fatty acid methyl esters (%) of the main fatty acids. Fatty acid methyl esters identification was based on retention times of authenticated commercial standards (AccuStandard and Sigma). Chromatographic data were processed by gas chromatography solution software (Supplier: JENCK S.A. Instrumental). Table 1 shows the dietary components and table 2 presents the fatty acid composition of each diet.

**Table 1.** Composition of diets.

| **Components** | **Diets** |  |  |
| --- | --- | --- | --- |
|  | **Control** | **Corn oil** | **Chia oil** |
| **Proteins** | 23% | 20% | 20% |
| **Fiber** | 6% | 5% | 5% |
| **Minerals** | 10% | 3.50% | 3.50% |
| **Calcium** | 1-1.4% | 1% | 1% |
| **Vitamins** | 2% | 1% | 1% |
| **Cistin/metionin/colin** | N/R | 0.55% | 0.55% |
| **Energy (Kjoule/G)** | 14% | 14% | 14% |
| **Startch** | N/R | 52.9% | 52.9% |
| **Carbohydrates** | 15% | 10% | 10% |
| **Lipid source** | 5% | 7% (corn oil) | 7% (chia oil) |

*This table shows the main composition of each diet. All diets were isoenergetic, providing 16.6 kJ/g. N/R: non reported. All diets were isoenergetic, providing 16.6 kJ/g.*

**Table 2.** Fatty acid composition of diets.

| Fatty acid | Diets |  |  |
| --- | --- | --- | --- |
|  | Control | Corn oil | Chia oil |
| **14:0 (myristic)** | 0.82 | 0.03 | Nd |
| **16:0 (palmitic)** | 17.54 | 12.21 | 6.83 |
| **16:1 n-9 (palmitoleic)** | 2.18 | 0.12 | Nd |
| **18:0 (stearic)** | 0.09 | 1.93 | 2.62 |
| **18:1 n-9 (oleic)** | 32.62 | 31.95 | 5.05 |
| **18:2 n-6 (linoleic)** | **35.55** | **51.26** | 18.91 |
| **18:3 n-3 (alpha-linolenic)** | 3.19 | 0.88 | **65.11** |
| **20:0 (arachidic)** | Nd | 0.50 | 0.17 |
| **20:1 n-9 (gondonic)** | Nd | 0.25 | Nd |

*This table shows the fatty acid composition of dietary oils (gas chromatography). Values are expressed as area percentage (a%) of total fatty acid methyl esters of the main fatty acids. The main fatty acid of each oil is highlighted.*

### Submandibular gland sample collection

30 days after starting the experimental diets, the animals were intraperitoneally anesthetized using phenobarbital (80 mg/kg body weight). Left and right submandibular glands were then removed and stored at -80 °C until processing. Finally, the animals were sacrificed by cervical dislocation.

### Endocannabinoid ligands

A fraction of glandular samples was lysed and 1 mL of supernatant was applied to a STRATA C-18 solid phase extraction cartridge (Phenomenex). Anandamide and 2-arachidonylglycerol were quantified by reverse phase high performance liquid chromatography (HPLC). A Phenosphere-Next C18 column (5 μm, 4.6 × 250 mm, Phenomenex) was employed. Metabolic separation was performed using a time program: a linear gradient from solvent A -methanol: water: acetic acid, 50: 50: 0.02 (v / v)-pH 6 to solvent B -methanol-for 20 min. A Beckman System Gold Model 166 programmable UV detector was linked to a computer for data processing. The values are expressed as total area percentage (a%).

### CB1 and CB2 receptors

These receptors were detected by indirect immunofluorescence. Samples were fixed in 4% formalin in PBS for 24 hours at room temperature, dehydrated, and embedded in paraffin. 5μm sections were collected on poly-L-lysine-coated glass slides and then incubated with monoclonal anti-CB1 and anti-CB2 antibodies (obtained from rabbit) diluted in TBS at 4 °C for 48 hours. After washing with PBS, slides were incubated with FITC-conjugated anti-rabbit IgG secondary antibodies (obtained from goat) at 37 °C for 1 hour [19], washed and observed through a Zeiss Axioskop 20 fluorescence microscope. Values are expressed as percentage of total positive marked area (%). Positive and negative controls were included in the study. Fiji ImageJ 1.53a software (NIH) was used for immunostaining quantification.

### Fatty acid amide hydrolase activity

Specific activity of this enzyme was determined by spectrophotometry (410 nm) with decanoyl p-nitroaniline [20] as substrate. Glandular samples were lysed in a solution containing 20 mM Hepes, 10% glycerol, 150 mM NaCl and 1% Triton X-100 at 4 °C, pH 7.8. After centrifuging, 20 µL of supernatant were taken and added to 175 µL of reaction buffer containing 1 µL of decanoyl p-nitroaniline fatty acid amide hydrolase substrate (Cayman Chemicals). The reaction was photocolorimetrically measured at 390 nm. Total protein concentration-employed to adjust the enzymatic activity-was determined by Bradford colorimetric technique. Values are expressed as μmol of product/μg of protein.

### Statistical analysis

The Kruskal Wallis test was applied to compare endogenous ligand levels, cannabinoid receptor immunostaining and fatty acid amide hydrolase activity between dietary groups, with a significance level of *p*≤0.05. Statistical Infostat program [21] was employed.

## Results

The n-6/n-3 ratios were significantly different between diets (Control=11.14; Corn oil =58.25; Chia oil=0.29) (*p*=0.001). Anandamide and 2-arachidonylglycerol were higher in Corn oil than in Control and Chia oil groups, but differences were statistically significant just for 2-arachidonylglycerol (*p*=0.003) (Fig.1). The greatest CB1 and the lowest CB2 receptor positively-marked area percentage were found in Corn (*p*=0.04) and in Chia oil groups (*p*=0.04), respectively (Fig. 2). Fatty acid amide hydrolase activity was higher in Corn than in Chia oil group (*p*=0.004) but not as compared to Control group (*p*=0.327) (Fig. 3).

**Figure 1.**
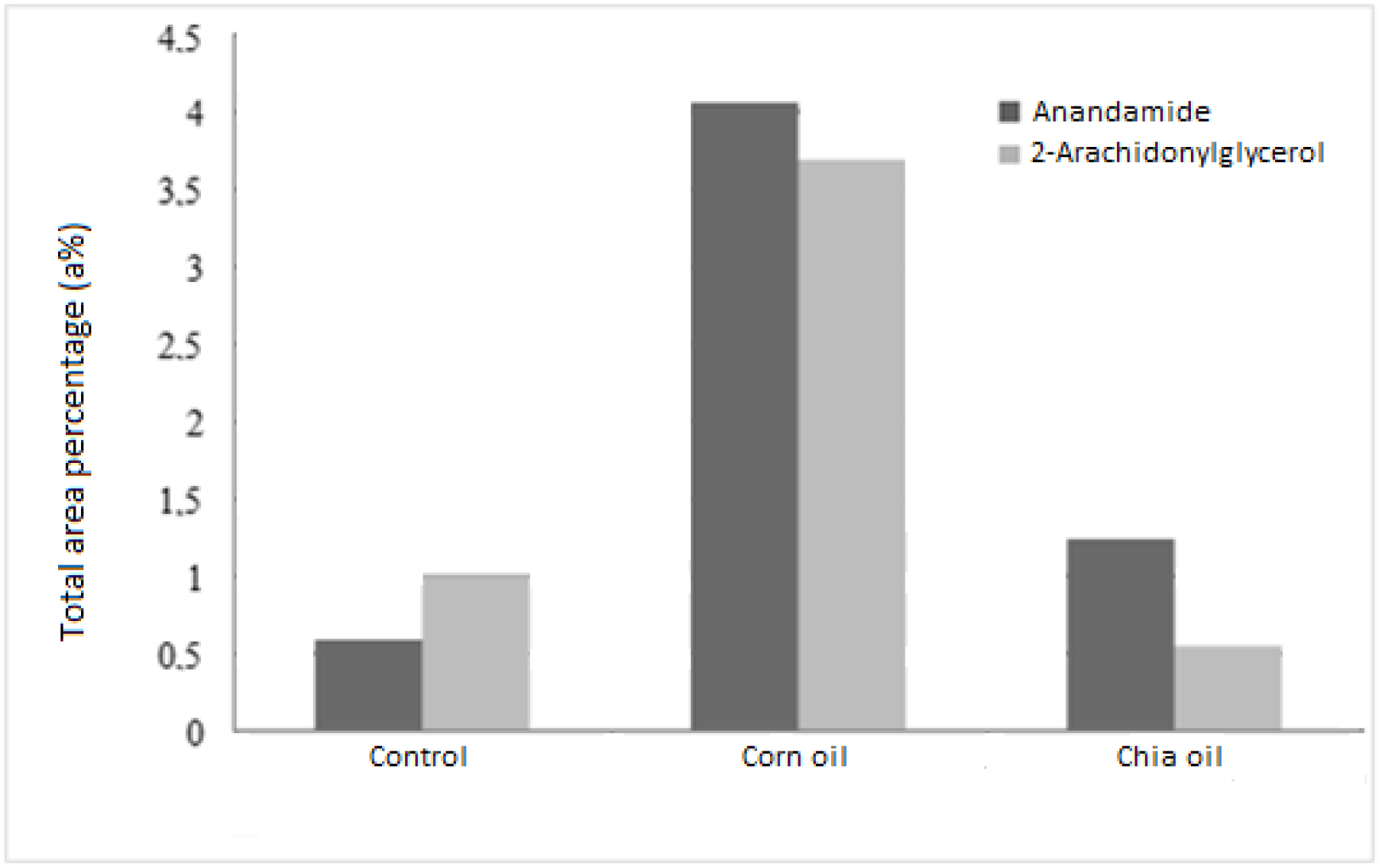
Endocannabinoid ligands. This figure shows the comparison between endocannabinoid ligands in each group. Values correspond to the mean of each group and are expressed as total area percentage (a%).

**Figure 2.**
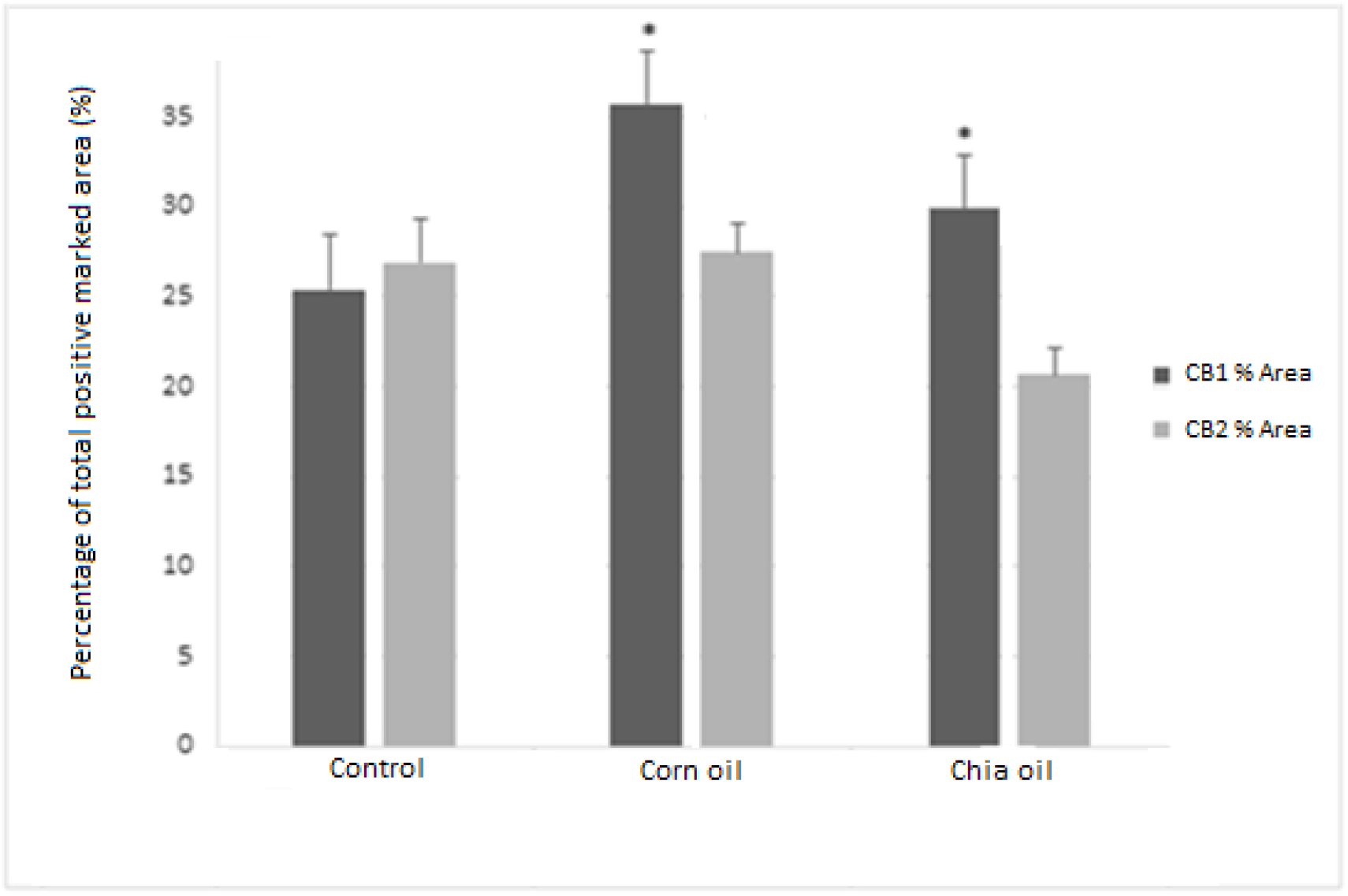
CB1 and CB2 receptors. This figure shows the comparison between cannabinoid receptors in each group. Values correspond to the mean of each group and are expressed as percentage of total positive marked area (%). * = Statistical significance.

**Figure 3.**
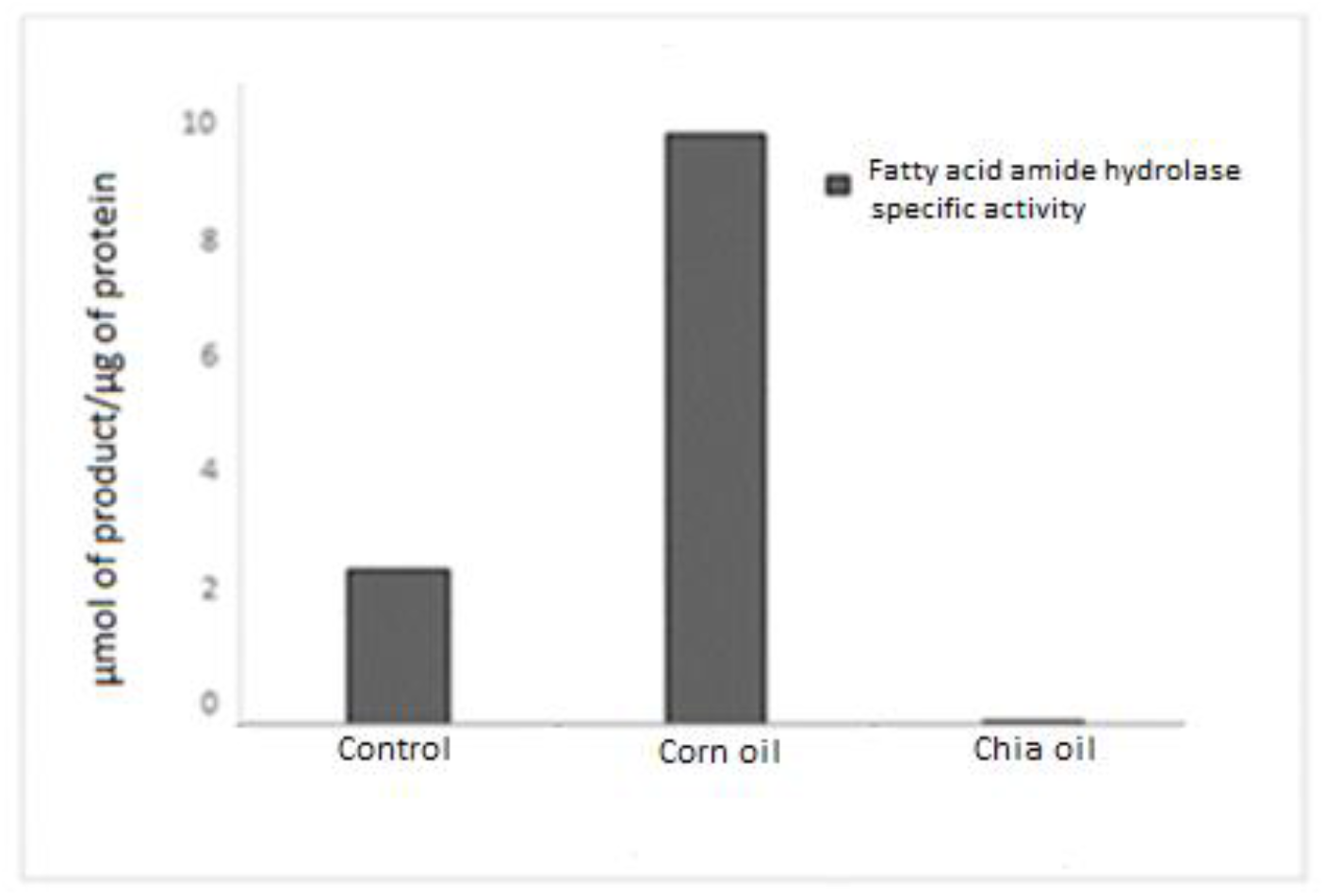
Fatty acid amide hydrolase activity. This figure shows the difference of specific activity of fatty acid amide hydrolase in each group. The values correspond to the mean of each group and are expressed as μmol of product/μg of protein.

## Discussion

The aim of the present study was to analyze the influence of dietary fatty acids on the endocannabinoid system in rat submandibular gland. Higher levels of anandamide and 2-arachidonylglycerol endogenous ligands were observed in Corn oil as compared to Control and Chia oil groups. Prestifilippo et al. [5] found that the injection of anandamide in the submandibular gland inhibits salivary secretion stimulated by norepinephrine and methacholine through the activation of CB1 and CB2 receptors, respectively. There are reports on the modulation of anandamide levels -in some organs and cell lines-by dietary fatty acid intake but no information could be found regarding submandibular glands [22]. However, in previous studies we have observed that arachidonic acid level -the main precursor of anandamide and 2-arachidonylglycerol-was lower in submandibular glands of animals fed on n-3 fatty acid-rich chia oil diet than in those fed on n-6 and n-9 fatty acid-rich corn and olive oil diets, respectively. In addition, a greater salivary flow was found in rats fed on n-3 fatty acid-rich chia oil diet for 30 days than in those which received n-6 and n-9 fatty acid-rich diets for 1 and 30 days, showing that salivary secretion could be modulated by dietary fatty acid intake and also by the time elapsed since their consumption [4].

Regarding cannabinoid receptors in submandibular gland, the highest expression of CB1 and the lowest of CB2 were observed in Corn and in Chia oil groups, respectively. It is known that the intake of a high fat diet as well as vitamin D may modulate the expression of CB1 receptors in the nervous system and in colon epithelium, respectively [23, 24, 25]. No references on the relationship between the expression of these cannabinoid receptors in submandibular glands and diet/dietary fatty acids could be found. However, other factors have shown to modify them. Pirino et al. [26] reported that the expression and localization of CB1 and CB2 in pig submandibular gland vary in response to masticatory activity (diets with different consistencies) suggesting that submandibular gland endocannabinoids modify salivary secretion, both qualitatively and quantitatively. Thoungseabyoun et al. [27] found that CB1 in rats is expressed mainly in the basolateral membranes of submandibular gland ductal cells but changes to the apical membrane of acinar and ductal cells when the salivary secretion is stimulated by isoprotererol.

It was observed that fatty acid amide hydrolase was also higher in n-6 fatty acid-rich Corn oil than in n-3 fatty acid-rich Chia oil dietary group. Touriño et al. [28] reported that levels of anandamide endocannabinoid ligand in different tissues involved in eating behavior and metabolism (hypothalamus, duodenum, jejunum and liver) were significantly higher in fatty acid amide hydrolase -/- mice as compared to wild-type animals, all of them fed on a high-fat diet. However, 2-arachidonylglycerol level was similar in both groups of mice. Engeli et al. [29] found a strong negative correlation between fatty acid amide hydrolase expression in adipose tissue of obese women and plasma levels of anandamide and 2-arachidonylglycerol. The results of this study show –for the first time according to our knowledge-that the habitual intake of diets rich in 18:3 n-3 alpha-linolenic and 18:2 n-6 linoleic acids modulates the endocannabinoid system components in rat submandibular gland. Particularly, a linoleic fatty acid-rich diet increases anandamide and 2-arachidonylglycerol ligands, CB1 and CB2 cannabinoid receptors and fatty acid amide hydrolase enzymatic activity. Although further research is required, it can be inferred –even on the basis of our previous study-that the intake of 18:2 n-6 linoleic acid decreases the salivary flow in rats, while the opposite occurs with n-3 fatty acid, due to the modulation of the endocannabinoid system.

Knowledge about the beneficial effects of n-3 fatty acids on salivary secretion could improve the quality of life of people with salivary secretion dysfunctions such as Sjögren’s syndrome and xerostomia related to medication, among others. Studies on salivary flow in relation to n-3 fatty acids incorporated through diet and/or nutritional supplementation should be carried out both in healthy subjects and in people suffering from salivary secretory dysfunction.

## Author contribution

The contribution of each author is described below:

**Conceptualization:** Jorge Darío Escandriolo Nackauzi, Adriana Beatriz Actis; **Methodology:** Gastón Repossi; **Validation:** César Nicolás Combina Herrera; **Formal analysis:** Jorge Darío Escandriolo Nackauzi, Gastón Repossi; **Investigation:** César Nicolás Combina Herrera, Mariana Blanco, Adriana Beatriz Actis; **Resources:** Adriana Beatriz Actis; **Data curation:** César Nicolás Combina Herrera, Mariana Blanco; **Writing – original draft preparation:** César Nicolás Combina Herrera, Adriana Beatriz Actis**; Writing – review and editing:** Mariana Blanco, Gastón Repossi, Jorge Darío Escandriolo Nackauzi, Adriana Beatriz Actis; **Supervision:** Jorge Darío Escandriolo Nackauzi; **Project administration:** Adriana Beatriz Actis; **Funding acquisition:** Adriana Beatriz Actis

